# Deciphering the CD73⁺ Regulatory γδ T Cell ecosystem associated with poor survival in Ovarian cancer

**DOI:** 10.1101/2024.12.09.627460

**Authors:** Ghita Chabab, Henri-Alexandre Michaud, Cécile Dejou, Laure-Agnès Chépeaux, Yaël Glasson, Florence Boissière, Marion Lenain, Anne-Sophie Dumé, Pauline Sarrant, Gabriel Chemin, Pauline Wajda, Bertrand Dubois, Anna MacManus, Pierre-Emmanuel Colombo, Michel Fabbro, Nathalie Bonnefoy, Maeva Chauvin, Virginie Lafont

## Abstract

The ability of tumor cells to overcome immune surveillance is an essential step in tumor development and progression. Among the immune cells playing a role in tumor control, γδ T cells contribute to the immune response against many tumor types through their direct cytotoxic activity against cancer cells and their capacity to regulate the functions of other immune cells. However, their presence in the tumor microenvironment is also associated with poor prognosis, suggesting that γδ T cells may also have pro-tumor activities. We previously described a regulatory γδ T-cell subset that expresses CD73 and produces IL-10, IL-8 and adenosine. Here, we report a higher CD73+ γδ T cell density in the tumor microenvironment of ovarian cancer samples from patients with short-term than long-term survival. Starting from this original observation, we investigated their neighborhood and described a specific ecosystem according to their pro-tumor functions.

## Introduction

Ovarian cancer is devastating due to its poor prognosis, with a 30% relative survival rate at 5 years, as it is usually diagnosed at an advanced stage (stage III/IV according to the FIGO system, which is a standardized system developed by the International Federation of Gynecology and Obstetrics) (1). Ovarian carcinoma accounts for over 90% of all ovarian cancer cases, and high-grade serous ovarian carcinoma (HGSOC) is the most prevalent and aggressive subtype, mainly due to resistance to platinum-based therapies (2,3). The mechanisms underlying recurrence, metastasis and resistance to therapy in ovarian cancer may be attributed partly to the tumor microenvironment (TME)which can suppress anti-tumor immunity by recruiting and promoting immune-suppressive cells, particularly from the immune compartment (4). In line with this, patients with HGSOC respond poorly to immune checkpoint inhibitors (5–7). The TME can affect, positively or negatively, solid tumor progression, including ovarian cancer (8). Indeed, the infiltration of various immune cell types into the TME is associated with the clinical outcome of patients with ovarian cancer. For example, tumor infiltration by CD3+ and CD8+ lymphocytes is associated with better survival in ovarian cancer (9,10), whereas the presence of infiltrating regulatory T cells (Tregs) predicts poor survival. Moreover, we previously showed that the density of γδ T cells is higher in ovarian tumor tissue than in normal matched ovarian tissue and tends to be higher in FIGO III/IV (i.e. advanced cancer) than low-stage tumors (11).

γδ T cells are unconventional T lymphocytes defined by the expression of a specific T cell receptor (TCR) consisting of the Vγ and Vδ chains. In humans, there are two main γδ T-cell subtypes, depending on the Vδ chain expressed: (i) Vδ2 T cells, mainly represented by the innate Vγ9Vδ2 T-cell subset, that predominate in blood and recognize phosphorylated non-peptide antigens, called phosphoantigens; and (ii) non-Vδ2 T cells, mainly represented by the Vδ1 subset, that predominate in tissues and recognize stress-induced antigens (12–14). Both subsets contribute to the immune response against cancer, particularly through their potent effector functions allowing direct elimination of tumor cells and the recruitment of other immune cells with anti-tumor activity (15). However, in addition to their anti-tumor functions, γδ T cells have also been associated with pro-tumor functions in some cancer types. Specifically, IL-17A-producing γδ T cells with pro-tumoral functions have been described in murine breast, ovarian and hepatocellular cancer models and in human colorectal cancer (16–18). In human breast cancer, γδ T cells have immunosuppressive functions associated with the induction of dendritic cell senescence (19). Furthermore, in murine and human pancreatic ductal adenocarcinoma, γδ T cells inhibit αβ T-cell activation and their infiltration via the PD-L1/PD-1 pathway, thereby favoring tumor progression (20). Similarly, it has been reported that in ovarian cancer, γδ T cells suppress the proliferation of CD4+ naive T cells (21). Altogether, these data support the hypothesis that some γδ T-cell subsets may have immunosuppressive functions and favor tumor progression in some solid tumor types. In agreement, we recently identified in human blood samples a γδ T-cell subset, found in both Vδ2 and Vδ1 populations, that expresses the ectonucleotidase CD73 and produces adenosine, an immunosuppressive molecule (22,23). This cell subset also produces IL-10 and IL-8 and inhibits conventional T-cell proliferation in a CD73/adenosine-dependent manner (22,23). Based on these findings, here, we explored the potential regulatory role of CD73+ regulatory γδ T cells in the immune landscape of ovarian cancer. Starting from our original observation that the presence of CD73+ γδ T cells is associated with poor clinical outcome in human ovarian cancer, we characterized the profile and investigated the neighborhood of both CD73+ and CD73-γδ T cell populations present in the ovarian TME and found a specific ecosystem according to their pro- and anti-tumor functions.

## Materials and methods

### Tissue sample collection

***Ovarian*** tissue samples were obtained from the Biological Resources Center of Montpellier Cancer Institute (ICM, France), and were collected from patients with HGSOC following the French regulations, under the supervision of the investigator Dr Fabbro. The collection was declared to the French Ministry of Higher Education and Research (declaration number DC-2008–695). The study was approved by the ICM Institutional Review Board (ICM-CORT-2020-32). Clinical data (e.g., age, treatment, TNM, grade) were obtained by reviewing the medical files. All samples were collected from biopsies taken at the time of diagnosis, prior to any treatment. Nearly all tumors (95% of patients) exhibited TP53 mutations. Due to the lack of variability in TP53 status, no meaningful stratification was performed based on this factor.

### Ovarian cancer tumor microarray (TMA)

TMAs regrouping 95 HGSOC samples were constructed to retrospectively compare long-term (LT, >5 years) vs short-term (ST, <5 years) ovarian cancer survivors. For each tumor sample, two cores (1 mm in diameter) were sampled from different malignant areas. Of the 95 tumors sampled on TMA, 91 (48 LT and 43 ST) could be analyzed, due to the absence of interpretable spots for 4 of them. The main clinicopathological characteristics of this cohort are presented in Table S1.

### Immunohistochemistry (IHC)

3 µm thin TMA sections were mounted on Flex microscope slides (Agilent Technologies) and dried at room temperature (RT) overnight before IHC. The PTLink system (Agilent Technologies) was used for de-paraffinization and heat-induced antigen retrieval in High pH Buffer (Agilent Technologies) at 95^◦^C for 15 min. IHC was performed using the Dako Autostainer Link48 platform (Agilent Technologies).

Briefly, after endogenous peroxidase neutralization, TMA sections were incubated with the mouse monoclonal anti-TCR δ antibody (Clone H-41, Santa Cruz Biotechnology) at RT for 30 min. This was followed by an amplification step with a mouse linker and the Flex Detection System (Agilent Technologies) and 3,3’-diaminobenzidine as substrate. Sections were counterstained with Flex Hematoxylin (Agilent Technologies). A NanoZoomer slide scanner system (Hamamatsu Photonics) was used to digitalize the stained TMA sections with a ×40 objective. Immunoreactive cells were identified on the digitalized slides with the NDP.view2 software.

### Immunofluorescence

After de-paraffinization, TMA sections underwent antigen retrieval using 1X Target Retrieval Solution (S2367, Agilent Technologies). Then, they were incubated in 1X Superblock Blocking Buffer (37515, Thermofisher) for 45 min followed by 1h incubation in FcR Blocking solution (130-059-901, Miltenyi Biotech). After washing, TMA sections were incubated with primary antibodies against γδ TCR (H-41, Santa Cruz, mouse, 1/25) and CD73 (D7AF9A, Cell Signaling Technology, rabbit, 1/100) at 4°C overnight and washed. Then, sections were incubated with secondary goat anti-rabbit Alexa Fluor Plus 555 (A32732, Thermofisher) and goat anti-mouse Alexa Fluor 647 (A21236, Thermofisher) antibodies. Finally, sections were counterstained with DAPI and imaged with an AxioScan (Zeiss) to obtain high-power field images. TMA sections were analyzed by two investigators blinded to the clinicopathological characteristics and survival outcomes. Cell density was assessed by counting the number of CD73- and CD73+ γδ T cells per spot (1 mm^2^) and calculated as the mean of two spots.

### Survival analysis

Categorical variables were presented as frequency distributions and continuous variables as medians and ranges. Categorical variables were compared with the Pearson’s chi-square test. Overall survival (OS) was defined as the time between the date of surgery and the date of death (whatever the cause). Patients lost to follow-up were censored at the last documented visit. The Kaplan–Meier method was used to estimate the OS. Differences in survival rates were compared using the log–rank test. Hazard ratios (HR) are given with their 95% confidence interval (95% CI). Statistical analyses were performed with STATA 16.0 (StatCorp, College Station, TX, USA).

### Tumor dissociation

Freshly samples of resected ovarian tumors from patients with HGSOC were cut into small pieces (∼1 mm^3^) and incubated in RPMI medium with a digestion solution (1mg/mL collagenase IV from Sigma - C5138-100mg - and 0.2 mg/mL DNase I from Roche - 11284932001) at 37°C for 30 min, interspaced by two rounds of mechanical dissociation with gentle flushing of the mixture. The obtained single-cell suspensions were filtered and stained for immunophenotyping, or incubated at 37°C with Golgi stop for 1 h in complete medium before performing immunostaining for cytokine analysis, as described in the next section.

### Flow cytometry analysis

After dissociation, cells were washed in cold PBS and incubated with a Fc receptor blocking reagent, to minimize non-specific binding of antibodies to Fc receptors (130-059-901, Miltenyi Biotec), and a fluorescent cell viability marker (C36628, Beckman Coulter) at 4°C for 15 min. Then, primary antibodies against surface markers were added in PBS-2% fetal calf serum (FCS) at 4°C for 30 min. If no intracellular staining was performed, cells were washed and fixed in 1% paraformaldehyde. If an intracellular staining was performed, cells were washed and then fixed and permeabilized using the FOXP3 Fixation and Permeabilization Kit (00-5523-00, eBioscience) at RT for 20 min, following the manufacturer’s protocol. Intracellular staining was performed by incubating cells with primary conjugated antibodies diluted in intracellular blocking solution at 4°C for 1 h. Cells were washed and fixed in 1% paraformaldehyde. The antibodies used for flow cytometry experiments are listed in Table S2. Data were acquired with a Cytoflex LX cytometer (Beckman-Coulter) and analyzed with the FlowJo software.

### Cancer-associated fibroblast (CAF) isolation from tumor samples and immunofluorescence staining

After tumor sample dissociation and filtration, the fraction left in the filters was collected and cultured in DMEM-F12 with 10% FCS until full adherence. The medium was changed every 2 or 3 days and cells were passaged once they reached confluence. Immunofluorescence staining with antibodies against alpha smooth muscle actin (α-SMA) (F3777, Sigma, 1/200) and vimentin (C9080, Sigma, 1/600) was used to confirm the CAF phenotype. After 15 days in culture and confirmation of their phenotype, CAFs were then used for ELISA experiments, as described below.

### Ovarian cancer cell line culture

Cells were cultured in DMEM-F12 with 10% FCS. Medium was changed every 2 or 3 days and cells were passaged once they reached confluence. Cells were then used for ELISA experiments as described below.

### IL-6 ELISA

1.10^5^ cells were plated in 48-well plates in 0.5 mL of complete cell culture medium (DMEM 10%SVF). After 24 h, supernatants were removed and 0.5 mL of fresh medium was added. IL-6 production in the supernatants of primary human ovarian CAF cells and four different human ovarian cancer cell lines (SKOV3, OVCAR3, OVCAR8, Kuramochi) was evaluated 24 h after changing the medium using the Human IL-6 DuoSet ELISA Kit (DY206-05, R&D Systems) following the manufacturer’s instructions.

### Imaging Mass Cytometry (IMC)-based tissue selection

Four ovarian tumors from the ST survivor group were selected based on the γδ T-cell infiltration density. From whole formalin-fixed paraffin-embedded tumor blocs, three sections were cut. One was stained with hematoxylin-eosin-saffron to identify tissue structures; one was used for IHC to localize γδ TCR, and one for IMC labeling and analysis. Based on the IHC data, 36 regions of interest (ROI) were selected and analyzed by IMC.

### IMC - tissue labeling

After de-paraffinization and antigen retrieval using the Dako Target Retrieval Solution at pH 9 (S2368, Agilent Technologies) in a water bath (96°C for 30 min), tissue sections were encircled with a Dako Pen, incubated with Superblock™ (37515, Thermofisher Scientific) at RT for 45 min, and then with human FcR Blocking Reagent (130-092-575, Miltenyi) at RT for 1 h. After three washes (8 min/each) in PBS/0.2% Triton X-100 (PBS-T), metal-tagged antibodies (list in Table S3) were diluted in PBS/1% BSA buffer. After incubation with the primary antibodies at 4°C overnight, sections were washed in PBS-T three times (8min/each) and nuclei were stained with iridium 191 and 193 (1:400 in PBS; Standard Biotools), a DNA intercalator, at RT for 30 min. Sections were washed in PBS for 5 min, then in distilled water for 5 min, and dried at RT for 30 min.

### IMC - data acquisition

Images were acquired with the Hyperion Imaging System coupled to a Helios time-of-flight mass cytometer (Standard Biotools) according to the manufacturer’s instructions. After selection, the ROIs were ablated with a UV laser at 200Hz. The Hyperion Imaging System was calibrated to ensure system stability and reproducibility following the manufacturer’s instructions. Data were exported as MCD files and visualized using MCD™ viewer 1.0.560.6.

### IMC – data processing and cell segmentation

Tiff files were extracted from the raw data, denoised and segmented using a custom pipeline based on the Steinbock framework with Docker container (Deepcell)(24). Then, the segmentation masks were extracted based on the iridium 191 and 193 nuclear staining and CD20, CD14, CD57, CD16, PanCK, CD3, CD66b, CD11c, CD4, CD68, TCR γδ, CD8a, CD15, GrB, Ki67, HLA-DR and PDL1 staining (membrane/cytoplasm markers). Default settings were kept for segmentation (pixel size at 1µm, whole-cell segmentation, no normalization, and mean aggregation). The mean signal intensities for each marker and for each cell were extracted from the .tiff files of each channel and the segmentation masks with the *computeFeatures* function of the R package EBImage (25). Then, data were exported in the fcs format with the R package flowCore (26).

### Manual gating

Hierarchically gating and specific marker intensity were used to identify immune cell subsets using FlowJo v10.10 (Supplementary Fig. 3A). Positivity thresholds were checked by comparing the signal intensity of a marker with its intensity on the raw image using an in-house application. Then, the gates were exported and reimported into R for further analysis.

### Single-cell clustering

Immune cell clustering was performed using the Rphenograph package with k=50 and based on normalized channels. Clusters with similar marker expression profiles were combined with cell types to account for over-clustering.

### IMC – Generation of neighbors

Neighboring cells were extracted with a homemade R function. To identify its neighbors, each cell was considered as the center of a circle. Then, the following formula derived from the Pythagorean theorem was used:

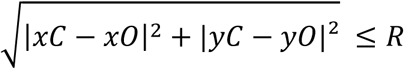

where *xC* and *yC* are the coordinates of the cell centroid of interest, *xO* and *yO* the coordinates of other cell centroids, and R the radius defined at 12µm. For each ROI, every cell for which the equation was true was considered a neighboring cell and exported in fcs format. The radius distance of 12µm was chosen to identify the cells that were most likely in contact with the cell of interest.

### Statistical analysis

The Mann-Whitney, log-rank, or Wilcoxon signed rank test was used to analyze the results depending on the experiment (the used tests are mentioned in the figure legends). A p value <0.05 was considered significant. Analyses were performed with GraphPad Prism, version 8. *p<0.05; **p<0.01; ***p<0.001; ****p<0.0001

## Results

### CD73+ γδ tumor-infiltrating lymphocyte (TIL) density correlate with worse clinical outcomes in patients with ovarian cancer

We and others showed that the density of γδ T cells is higher in ovarian cancer than in normal ovarian tissue samples and that the number of γδ T cells is positively correlated with advanced clinicopathological features of ovarian cancer (11). However, the potential role of γδ T cells in ovarian cancer outcomes remains unclear. A study reported that in ovarian cancer, outcome is positively influenced by the functionality of γδ T cells, measured by their ability to produce Th1 cytokines, and negatively affected by the proportion of CD39+ T cells (27). Therefore, to clarify their role, we analyzed the impact of γδ T-cell density on cancer progression in a cohort of 91 patients with ovarian cancer (n=43 ST and n=48 LT survivors) for whom clinical data were collected over the years. Regardless of whether patients received chemotherapy before or after surgery, all analyzed samples were collected before treatment. Their median follow-up was 13.44 years (95% CI [11.67-15.47]) and their median survival was 5.14 years (95% CI [3.75 - 6.08]). We first quantified by immunofluorescence staining all γδ T cells and then the CD73- and CD73+ γδ T-cell subtypes. The densities of total (Fig. 1A), CD73+ (Fig. 1B) and CD73- (Fig. 1C) γδ T cells were significantly higher in ST survivors than in LT survivors, suggesting that the presence of γδ T cells at the tumor site is associated with poor survival. We next investigated the association between total, CD73+ and CD73-γδ T cells and OS in these 91 patients. Overall survival (OS) was significantly longer in patients with low density of γδ T cells compared to those with a high density (p < 0.05; median OS [IC50] low density = 6.099 years vs. high density = 3.685 years; Fig. 1D). Similarly, OS was significantly shorter in patients with high density than those with low density of CD73+ γδ T cells (p < 0.001; OS [IC50] low density = 6.077 years vs. high density = 3.229 years; Fig. 1E) and this difference is even more pronounced than that observed for total γδ T cells. The OS [IC50] of patients with high CD73⁺ γδ T cell density was even shorter than those with high total γδ T cell density (3.229 years vs 3.685 years), suggesting a more detrimental role of CD73⁺ γδ T cell subset. In contrast, CD73⁻ γδ T cell density did not significantly affect survival although OS was lower for high density (p = ns; OS [IC50] low density = 6.099 years vs high density = 4.175 years; Fig. 1F). These reinforce the hypothesis that although the density of γδ T cell subsets is higher in short-term (ST) survivors, the effect γδ T cells on survival is mostly due to the CD73+ subset suggesting that the presence and density of CD73+ γδ T cells may be a potential prognostic marker in ovarian cancer.

**Figure 1.**
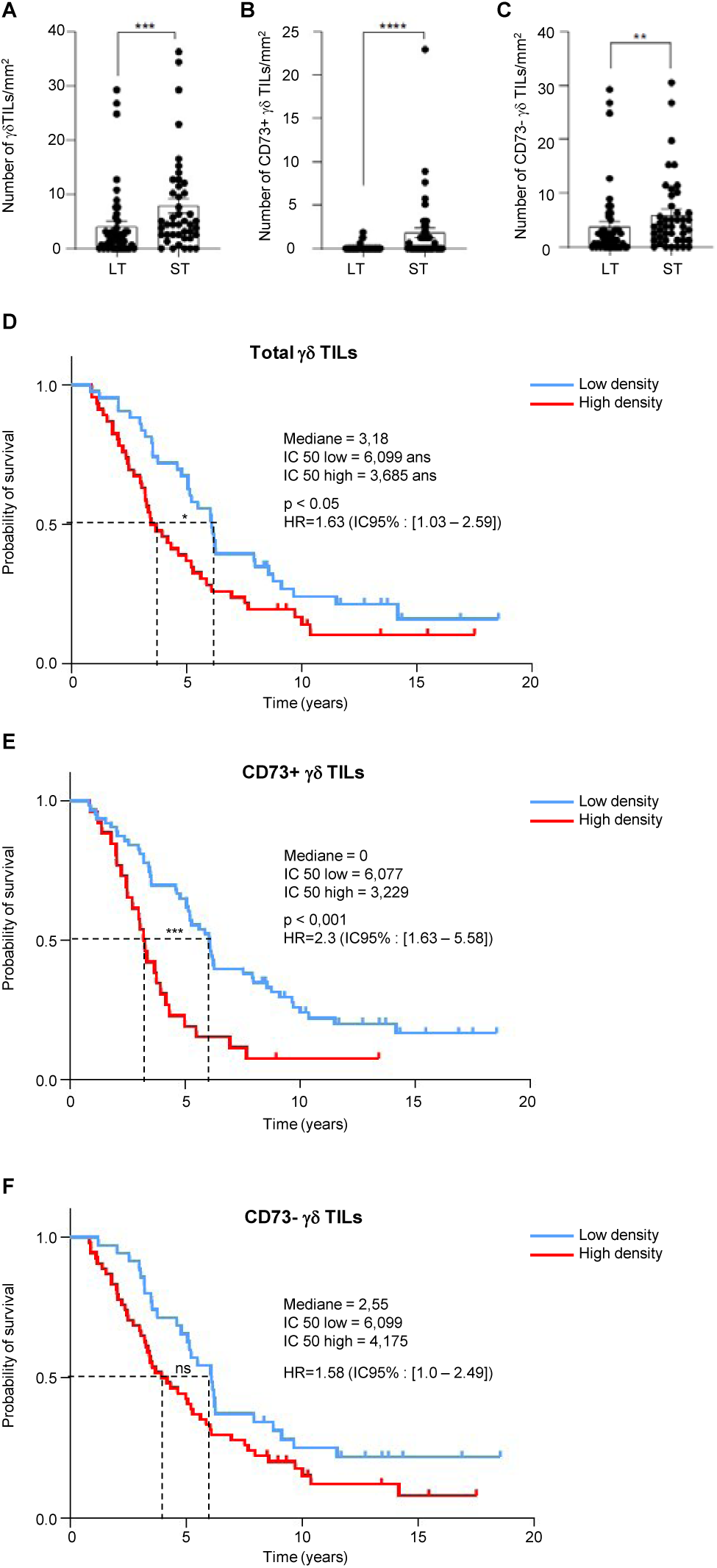
Relationship between γδ T cell infiltration and survival outcomes in ovarian cancer patients or correlation between γδ T cell infiltration and survival in patients with ovarian cancer. Scatter plots comparing the density of total γδ T cells (A), CD73+ γδ T cells (B), and CD73-γδ T cells (C) in short-term (ST, n=43) and long-term (LT, n=48) survivors. Overall survival analysis according to the density of total γδ T cells (D), CD73+ γδ T cells (E), and CD73-γδ T cells (F) in the same patients with ovarian cancer. Data are the mean ± SEM. ns: not significant; **p < 0.01; ***p < 0.001; ****p < 0.0001 (Mann-Whitney test for panels A, B and C; log-rank test for the survival analysis). TILs, tumor-infiltrating lymphocytes.

In parallel, the analysis of TCGA data revealed a strong association between a CD73+ tumor microenvironment and poor patient outcomes (Fig. 2A). To investigate whether this CD73+ microenvironment correlates with γδ T cell infiltration, contributing to reduce survival as demonstrated in Figure 1, we analyzed single-cell RNA sequencing (scRNA-seq) data from 20 untreated ovarian cancer samples. These samples were extracted from the GSE147082, GSE241221, and GSE235931 datasets and pooled with Seurat’s integration pipeline (28,29) (Fig. 2B). Patients were divided into two groups based on median NT5E gene expression levels (gene encoding CD73): CD73/NT5E-Low and CD73/NT5E-High (n=10 per group) (Fig. 2B). Interestingly, expressions of specific γδ T cell transcripts such as TRDC (coding the TCRδ chain), TRGC1 and TRGC2 (both coding TCRγ chain) are higher in TME overexpressing NT5E (CD73) (Fig 2C). In addition, the following cytokine expression (CXCL8, IL-10, IL-6) (Fig. 2D) was higher in the CD73/NT5E-High than CD73/NT5E-Low group as well as higher proportion of CAFs and γδ T cells (Fig. 2E). These population enrichments suggest that the expression of IL-6, IL-10 and CXCL8 and the density of γδ T cells and CAFs are significantly higher in CD73/NT5E-rich TMEs than in CD73/NT5E-low TMEs. To validate these observations and to further explore these relationships between cell subsets in different TMEs, we performed a spatial proteomic analysis using an innovative high-plexed technology that allows detailed profiling of cell phenotypes and the tumor ecosystem.

**Figure 2.**
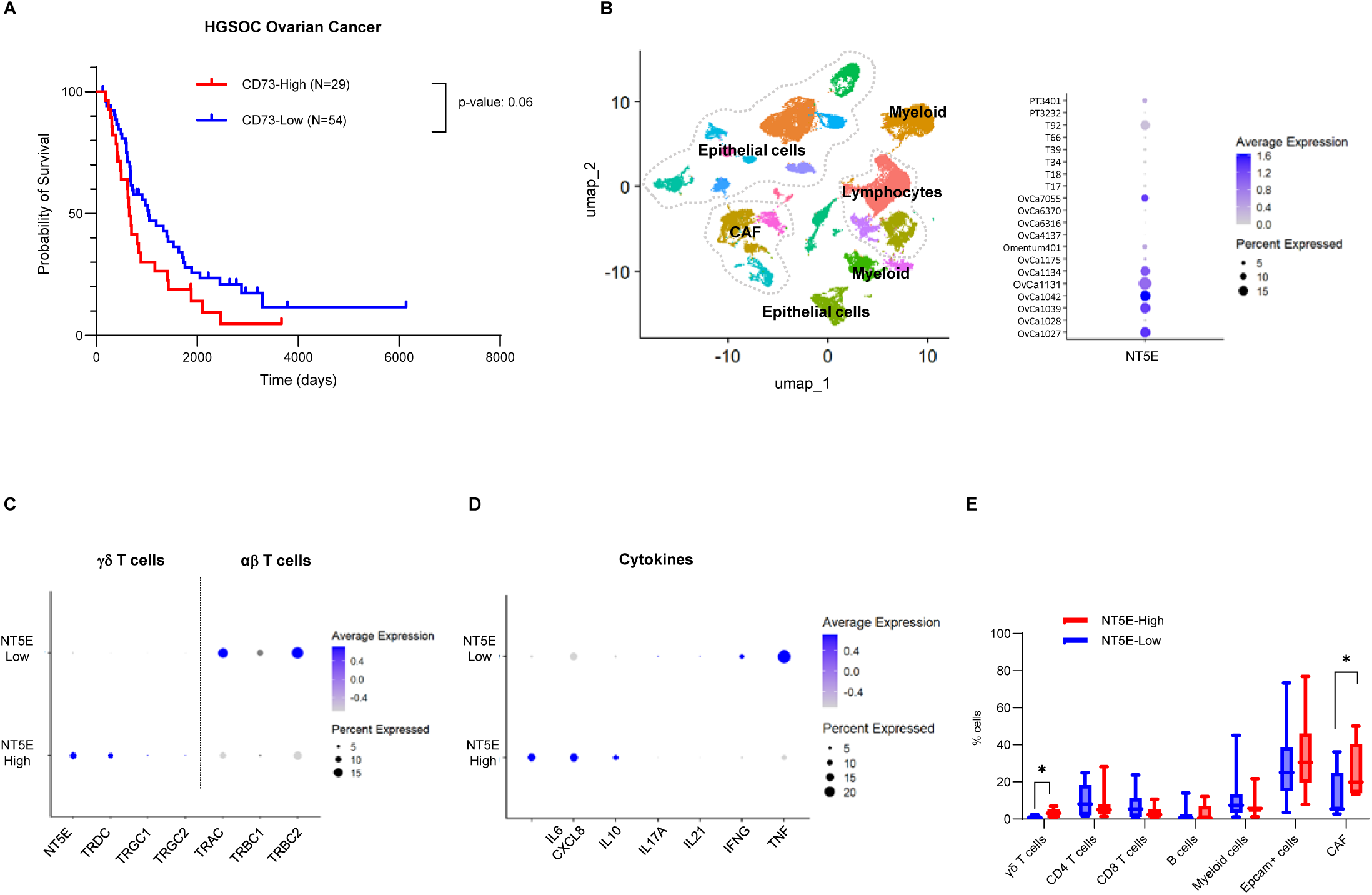
Analysis of single-cell RNA sequencing data from 20 naive HGSOC patients. (A) Kaplan-Meier survival plots comparing patient populations with low and high CD73 (NT5E) transcriptomic expression according to the TCGA dataset stratified on the median expression. (B) UMAP plot of single-cell RNA-sequencing data from 20 untreated high-grade serous ovarian cancer (HGSOC) patients, integrating publicly available datasets GSE147082, GSE241221, and GSE235931. Tumor-associated cell populations were identified based on transcriptomic clustering and annotated accordingly. Right panel: Dot plot illustrating the expression of NT5E (CD73) across the 20 patients, aggregated at the whole-tumor level. Dot size indicates the proportion of cells expressing NT5E within each cluster; color intensity reflects the average expression level. (C) Dot plot comparing the expression of T cell receptor (TCR) chain genes between patients with low (NT5E_Low, n = 10) and high (NT5E_High, n = 10) NT5E expression, defined by a median split of average NT5E transcript levels across all patients. Dot size represents the proportion of cells expressing each gene; color intensity indicates the average expression level. (D) Dot plot showing the expression of cytokine genes in the NT5E_low and NT5E_high patient groups, using the same visual scale as in panel C. (E) Bar plot displaying the relative proportions (%) of major cell types within the tumor microenvironment, stratified by NT5E expression group, based on single-cell cluster annotation.

### Cell composition of human ovarian tumors from short-term survivors

First, we investigated the tissue composition of ovarian tumor samples of 4 patients from the ST survivor group of our cohort by multiplexed IMC using a specific protocol described in Supplementary Fig. 2. We selected 38 region of interests (ROIs) corresponding to approximately 56 mm^2^ based on the γδ T-cell density identified by IHC, on a serial section. Using a 31-marker panel (Table S3) and a specific gating strategy to identify tumor, stromal and immune cells (Supplementary Fig. 3A-B), we analyzed the cell composition of the selected ROIs (Supplementary Fig. 3C-D). We identified 17 cell types and observed heterogeneity among tumor samples and ROIs. Tumor cells, identified as E-cadherin+ and PanCK+ cells, were the most important cell type in all samples (46% of all cells), followed by immune cells (27% of all cells) and fibroblasts (vimentin+ cells; 15% of all cells) (Supplementary Fig. 3D-E). Moreover, α-SMA+ cells were 8% of all cells and <1% of cells represented the vascular compartment (Supplementary Fig. 3E). The remaining cells (<3.5%), classified as “undetermined” (Supplementary Fig. 3E), corresponded to cells with a nucleus, but that could not be identified using our panel. The characterization of the immune infiltrate (Supplementary Fig. 3D and F) showed that lymphoid cells represented ∼75% of the immune infiltrate and most of them were T cells. In the myeloid compartment, monocyte-derived dendritic cells were the most abundant subset (∼13%).

### Phenotypic analysis of CD73+ and CD73-*γδ* TILs

Based on γδ TCR expression, we analyzed the γδ T-cell compartments by IMC (Fig. 3A). The proportion of γδ T cells was quite homogenous in all ROIs. They represented ∼3% of the immune compartment (Fig. C and ∼20% of all γδ TILs were CD73+ (Fig. 3D). Although the ROIs analyzed by IMC were selected based on the presence of γδ T cells, the results were consistent with the flow cytometry (FC) data obtained using fresh ovarian tumor samples (n=7) (Fig. 3B-D).

**Figure 3.**
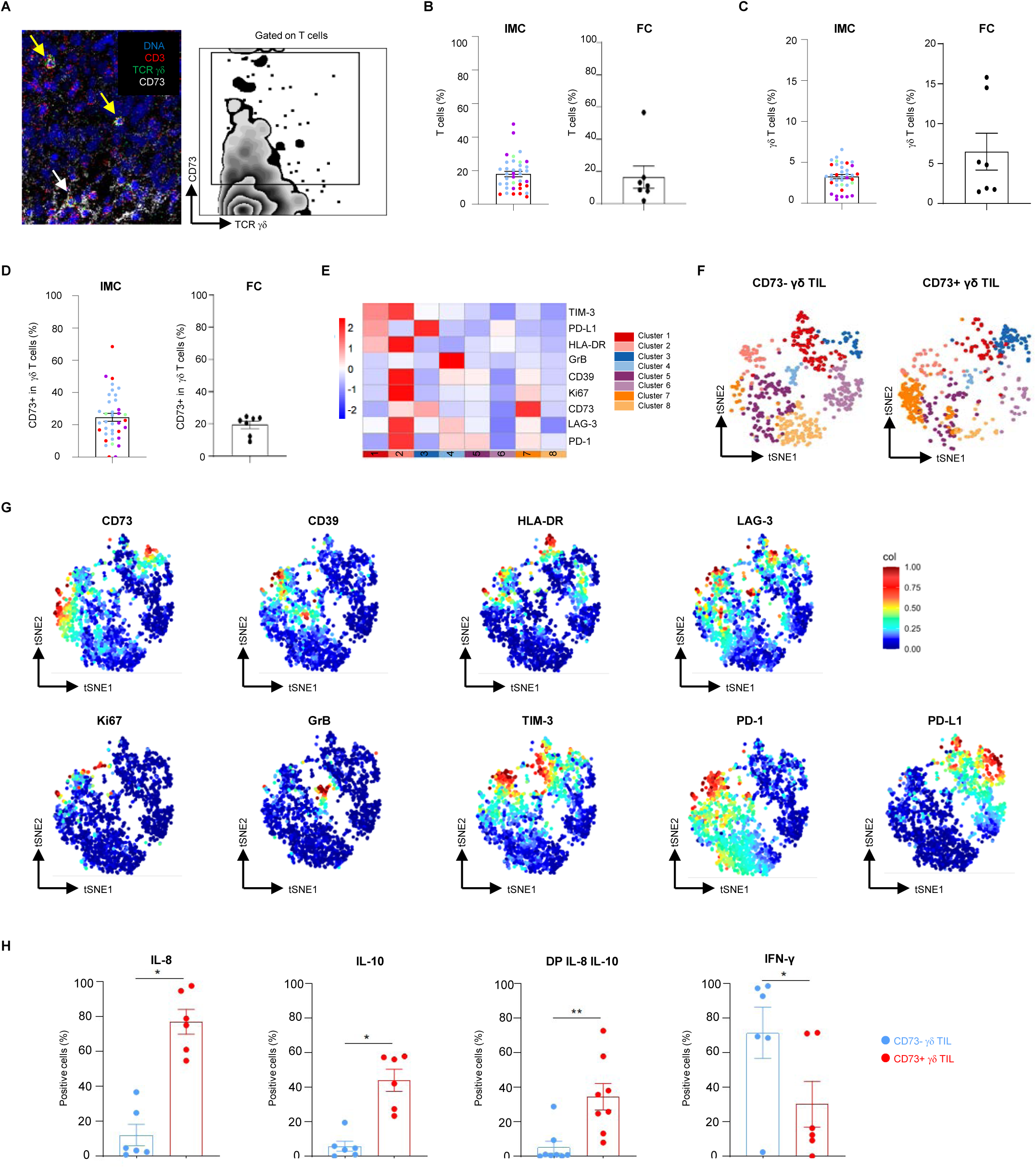
γδ T cell phenotype and function in ovarian cancer tissue. (A) IMC data showing CD73+ and CD73-γδ T cells in an ovarian tumor section (left panel) and the gating (right panel). Dot plots show the percentages of T cells (B), γδ T cells (C), and CD73+ γδ T cells (D) obtained through IMC analysis of 38 ROIs from four patients and flow cytometry analysis of seven patients. For IMC plots, each circle represents a ROI, and each color corresponding to an individual patient. For flow cytometry, each black circle represents an individual patient sample. (E) Heatmap of the mean of signal intensity of each marker and showing the clustering of γδ based on the expression of the listed markers. (F) tSNE plots showing cluster repartition in CD73+ and CD73-γδ T cells. (G) tSNE plots showing the expression of each analyzed marker in the total γδ T cell clusters. (H) Dot plots showing the percentage of CD73+ (red) and CD73-(blue) γδ TILs expressing IL-8, IL-10, double positive IL-8/IL-10 and IFN-γ. Each dot represents an individual patient. Data are the mean ± SEM (n=6); **p < 0.01; *p < 0.05 (Wilcoxon signed-rank test).

Unsupervised analysis of total γδ T cells from the IMC data allowed identifying eight clusters based on the expression of immune checkpoints and functional markers (Fig. 3E). The tSNE plots of the CD73+ and CD73-γδ T-cell subsets were clearly different (Fig. 3F). Clusters 1, 5, 6 and 8 were more abundant in the CD73-γδ T-cell subset, whereas clusters 3 and 7 were more abundant in the CD73+ γδ T-cell subset (Fig. 3F and Supplementary Fig. 4A). Cluster 3 was characterized by strong expression of PD-L1 and to a lesser extent of LAG-3 and TIM-3 (Fig. 3E and G) and cluster 7 by expression of PD-1, LAG-3, CD39 and Ki-67. Among the most abundant clusters in the CD73-γδ T-cell subset, cluster 1 was characterized by high expression of the functional marker HLA-DR and the immune checkpoints TIM-3, PD-L1; cluster 5 by expression of the immune checkpoints PD-1, CD39, LAG-3, and cluster 6 by expression of HLA-DR and PD-L1. Cluster 8 was not characterized by any of the analyzed markers (Fig. 3E-G). Clusters 2 and 4 were not specific to any subset. In parallel, we performed conventional flow cytometry analysis of immune checkpoint marker expression in fresh ovarian tumor samples (n=7). We confirmed the higher expression of PD-L1 in the CD73+ γδ T-cell subset. PD-L2 also was expressed in this subset, whereas TIGIT was more strongly expressed in the CD73-γδ T-cell subset (SupplementaryFig. 4B). This suggested that PD-L1 or PD-L2 when expressed may also contribute to the immunosuppressive functions of CD73+ γδ T cells.

We previously showed that regulatory γδ T-cell subsets, derived from *in vitro* culture of blood healthy donors, are characterized by CD73 expression and IL-8 and IL-10 production. Therefore, we investigated by flow cytometry the expression IFN-γ, IL-10 and IL-8 by γδ T cells isolated from fresh ovarian cancer samples in correlation with CD73 expression (Fig. 3H and supplementary Fig. 4C). The frequency of IL-8+ and IL-10+ cells was significantly higher in the CD73+ γδ than in the CD73-γδ T-cell subset (75% versus 10% for IL-8, 45% versus 5% for IL-10). Moreover, 35% of CD73+ γδ T cells expressed both cytokines. Conversely, the percentage of IFN-γ+ cells was significantly higher in the CD73-than CD73+ γδ T-cell subset (70% versus 30%). Altogether, these results demonstrated that CD73+ γδ T cells from ovarian tumors have a profile similar to that of the previously described regulatory CD73+ γδ T-cell subset (22,23) and suggested that their immunosuppressive properties may be involved in the poorer clinical outcome.

### CD73+ and CD73-*γδ* TILs have phenotypically and functionally different ecosystems

Besides the immune cell composition, the spatial location and interactions of immune cells within the TME are essential for their functions and the effect on tumor development and response to treatment.

Here, we investigated the γδ T-cell ecosystem by comparing the effector (CD73-) and regulatory (CD73+) γδ T-cell neighborhoods and interactions by IMC. After identification and localization of CD73+ and CD73-γδ T cells in ovarian cancer tissue sections (Fig. 4A), we measured and compared the individual interactions between CD73+ and CD73-γδ T cells with the other identified cell types. For both cell subtypes, we observed an heterogenous environment but with significant differences. CD73-γδ T cells had a neighborhood significantly enriched in tumor cells (24.4% versus 12.7% for CD73+ γδ T cells) (Fig. 4B and Supplementary Fig. 5). Conversely, fibroblasts and CD4+ T cells tended to be more frequently in contact with CD73+ than CD73-γδ T cells (Fig. 4B and C and Supplementary Fig. 5), suggesting that the CD73+ γδ T-cell neighborhood is enriched in fibroblasts and CD4+ T cells. Comparison of the CD73+ γδ T-cell ecosystem with the conventional regulatory T-cell (CD3+CD4+FOXP3+) ecosystem revealed a specific CD73+ γδ T cell ecosystem (Supplementary Data Fig6).

**Figure 4.**
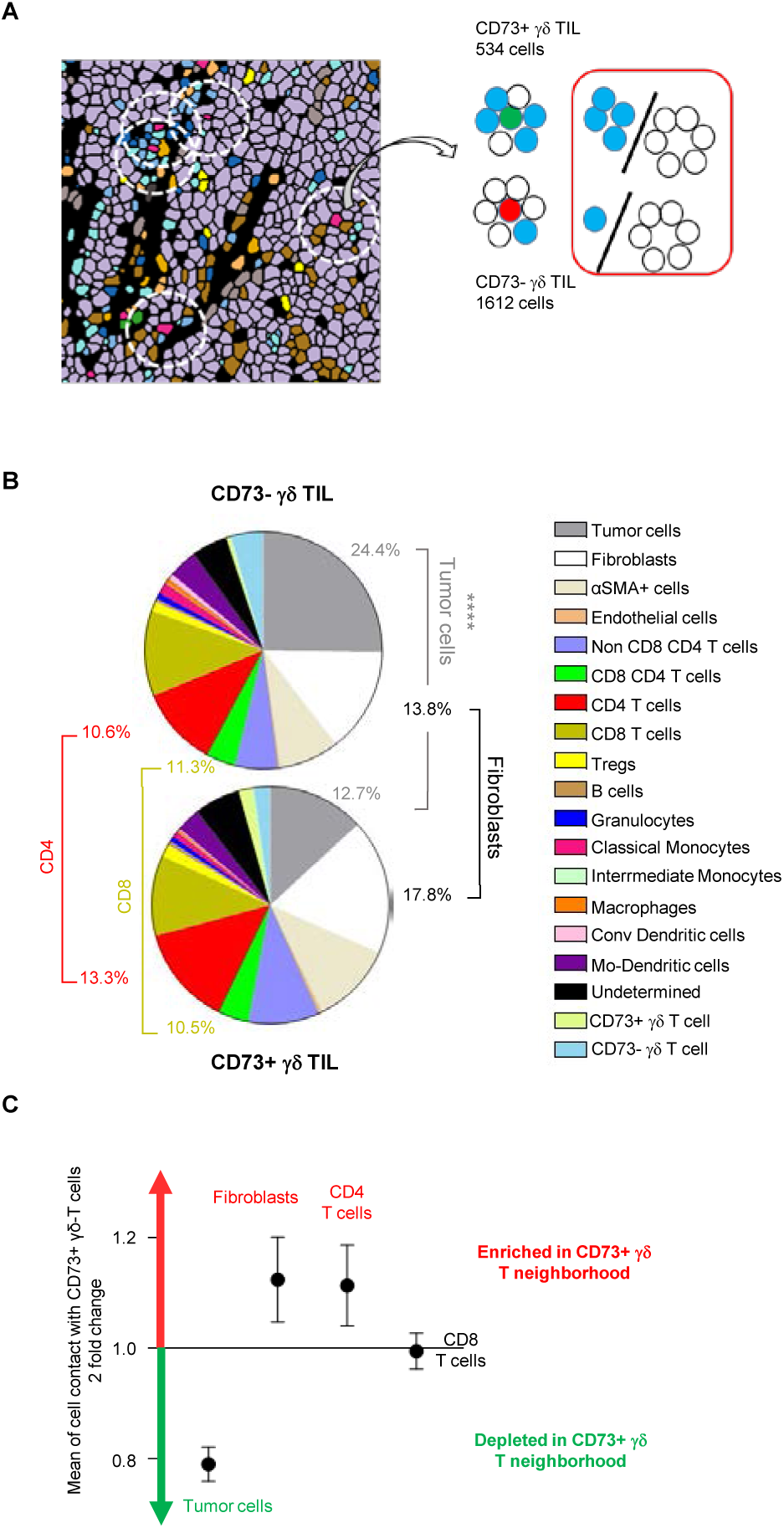
Neighbors of CD73+ and CD73-γδ T cells in ovarian tumor. (A) IMC strategy to extract all neighbors of the CD73+ and CD73-γδ T-cell subsets. (B) Pie charts showing the mean value of each identified cell type in the selected ROIs. Upper panel: Neighbors of CD73-γδ T cells; lower panel: Neighbors of CD73+ γδ T cells. ****p <0.0001 (Wilcoxon signed rank test). (C) The neighborhood cell enrichment represented as the log2 fold change of cell frequency in contact with CD73+ γδ T or CD73-γδ T cells.

We then determined the number of cells for all identified populations and chose to perform a functional analysis only of the cell compartments containing more than 750 cells to obtain robust results (Supplementary Fig. 7): tumor cells, fibroblasts, CD4+ and CD8+ T cells.

Unsupervised clustering of the tumor cell compartment allowed identifying eight cell populations that were differentially abundant depending on whether they were in the vicinity of effector (CD73-) or regulatory (CD73+) γδ T cells (Fig. 5A and Supplementary Fig. 8A). Indeed, tumor cells expressed CD73 and the epithelial-to-mesenchymal transition (EMT) marker ZEB-1, in the neighborhood of CD73+ γδ T suggesting migratory capacity (Fig. 5A). Tumor cells in contact with CD73-γδ T cells expressed PD-L1. Altogether, these data suggest that tumor cells in contact with CD73+ γδ T cells have a more aggressive phenotype. Similarly, cluster analysis showed that fibroblasts in contact with CD73+ γδ T cells expressed CD73 and PD-L1, whereas those in contact with CD73-γδ T cells expressed more CD39 and Ki-67 (Fig. 5B and Supplementary Fig 8B).

**Figure 5.**
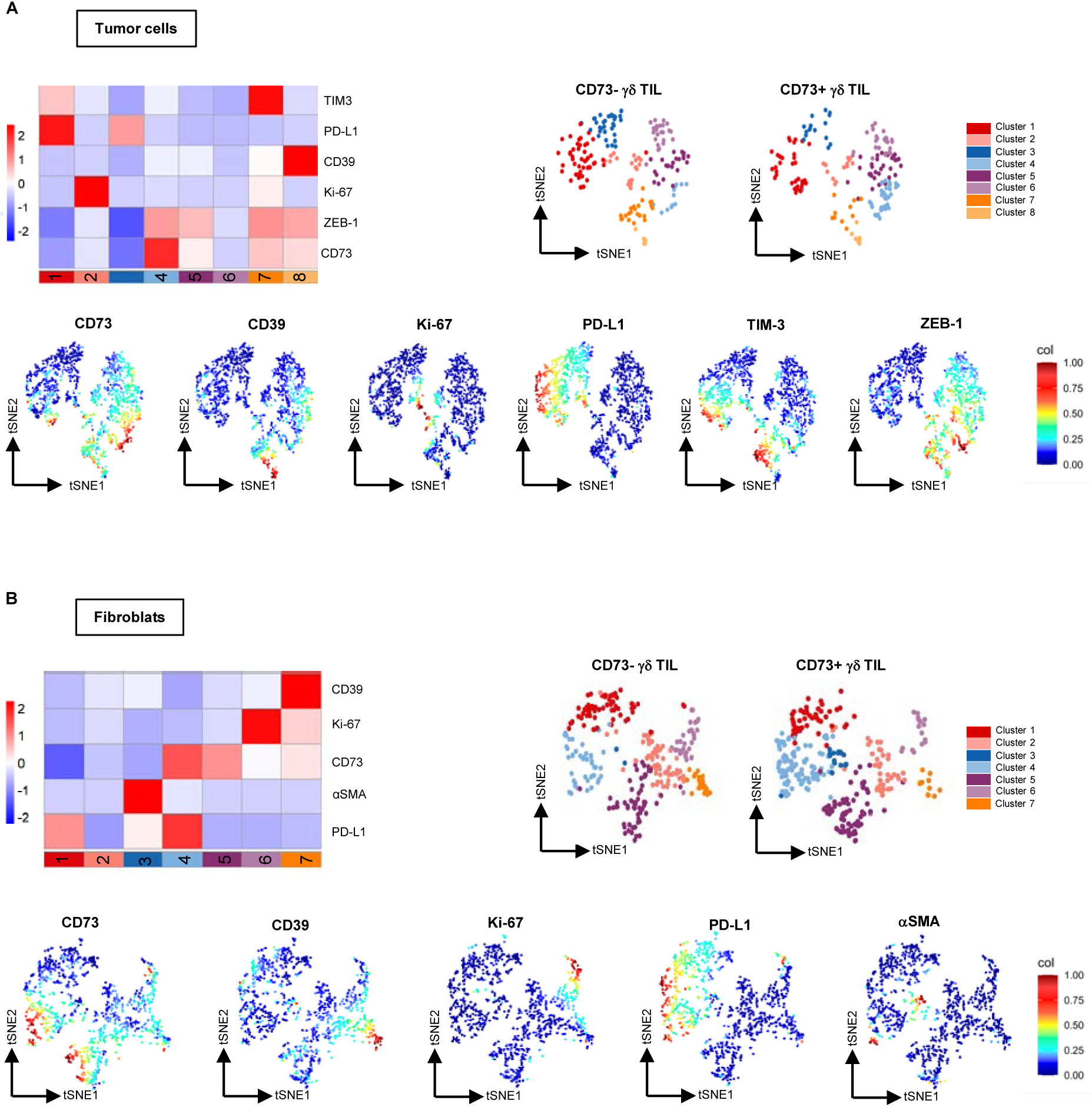
Phenotype of stromal neighbors of CD73+ and CD73-γδ T cell subsets. (A) Heatmap of the mean of signal intensity of each marker and showing the clustering of tumor cells based on the expression of the listed markers (upper left panel) and tSNE plots showing the tumor cell cluster repartition in CD73+ and CD73-γδ T cells (upper right panels). tSNE plot representation of normalized single marker expression in all the sin the total tumor cell clusters (lower panels). (B) Heat map showing the clustering of fibroblasts based on the expression of the listed markers (upper left panel) and tSNE plots showing cluster repartition in CD73+ and CD73-γδ T cells (upper right panels). tSNE plots showing the expression of each analyzed marker in the total tumor cell clusters (lower panel).

Like for other cell types, unsupervised analysis of the T-cell compartment gave different tSNE clustering plots in the neighborhood of CD73+ and CD73-γδ T cells for the CD4+ and CD8+ T cell populations (Fig. 6A-B and Supplementary Fig. 8C-D). The cluster analysis revealed that CD4+ T cells in contact with effector γδ T cells (i.e. the CD73-subset) express HLA-DR and/or PD-1, both activation markers. In contrast, cells that interacted with CD73+ γδ T cells displayed expression of CD73, PD-1, CD39, and LAG-3, which may indicative of an exhausted phenotype (Fig. 6A). For CD8+ T cells, the CD73+ and PD-1+ cluster was much more abundant in the neighborhood of CD73+ γδ T cells (Supplementary Fig 8D).

**Figure 6.**
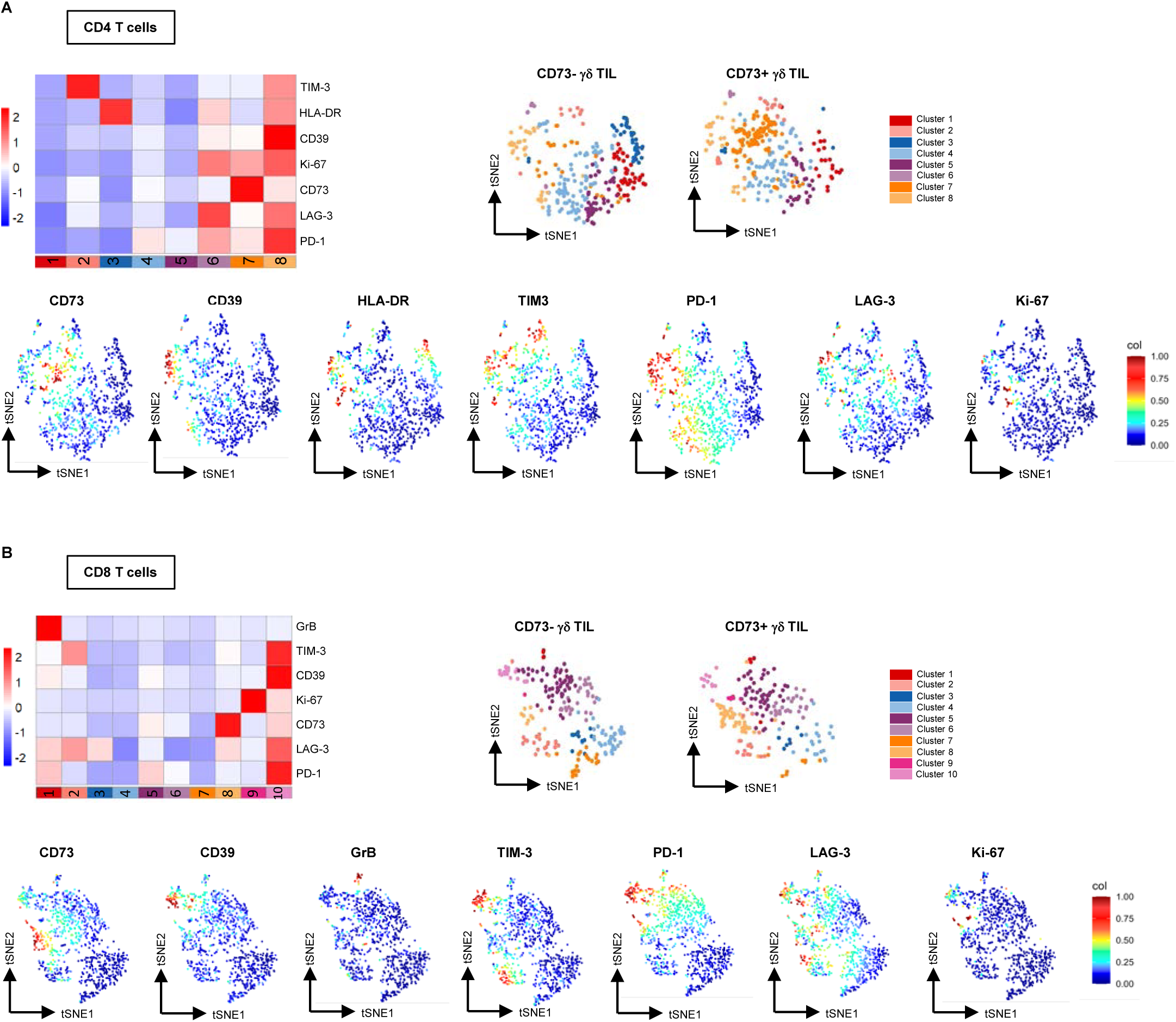
Phenotype of T cell neighbors of CD73+ and CD73-γδ T cell subsets. (A) Heat map showing the clustering of CD4+ T cells based on the expression of the listed markers (upper left panel) and tSNE plots showing cluster repartition in CD73+ and CD73-γδ T cells (upper right panels). tSNE plots showing the expression of each analyzed marker in the total tumor cell clusters (lower panels). (B) Heat map showing the clustering of CD8+ T cells based on the expression of the listed markers (upper left panel) and tSNE plots showing cluster repartition in CD73+ and CD73-γδ T cells (upper right panels). tSNE plots showing the expression of each analyzed marker in the total tumor cell clusters (lower panels).

Although the density of fibroblasts and T cells surrounding CD73- and CD73+ γδ T cells was similar, the functional phenotype of these cell populations differed, resulting in distinct ecosystems, suggesting that CD73+ γδ T cells are located in an immunosuppressive niche in close contact with tumor cells. Comparison of CD73+ γδ T cells and Treg revealed specific neighborhoods for each population (Supplementary Fig. 6), suggesting that the mechanisms responsible for establishing an immunosuppressive microenvironment are specific to CD73+ regulatory γδ T cells.

### Impact of IL-6 produced by ovarian CAFs and tumor cells on *γδ* T cells

We and others provided evidence that soluble factors, such as cytokines, can modulate the functions of γδ T cells (30). Specifically, we showed that IL-21 promotes the development of a pro-tumoral γδ T-cell subset, characterized by CD73 expression and production of immunosuppressive molecules, such as adenosine, IL-10 and IL-8 (22). Others have shown that CAFs from breast tumors produce IL-6 and induce the differentiation of CD73+ γδ T cells from normal breast tissue into immunosuppressive T cells (31). As IL-6 is strongly expressed in the ovarian TME (32), we hypothesized that CAF-produced IL-6 may be involved in the development or differentiation of CD73+ γδ T cells in ovarian cancer. We isolated CAFs from freshly resected ovarian tumor samples (n=3) and confirmed their phenotype/purity by α-SMA and vimentin staining (Fig. 7A). We then measured IL-6 in their supernatants and in the supernatants of four ovarian tumor cell lines by ELISA. Both CAFs and ovarian tumor cell lines produced IL-6, but the amounts were much higher in CAF supernatants than in tumor cells (1500 pg/mL vs ∼70pg/mL) (Fig. 7B). Additionally, as reported by other teams and by our group (11,22,31), we confirmed that IL-6, similar to IL-21, promotes the development of the CD73+ γδ T cell subset from blood-derived γδ T cells of healthy donors. (Fig. 7C and D). Altogether, these data suggest that the cross-talk between CAFs and γδ T cells in the TME through IL-6 production may favor an immunosuppressive niche that promotes tumor growth and immune escape.

**Fig. 7.**
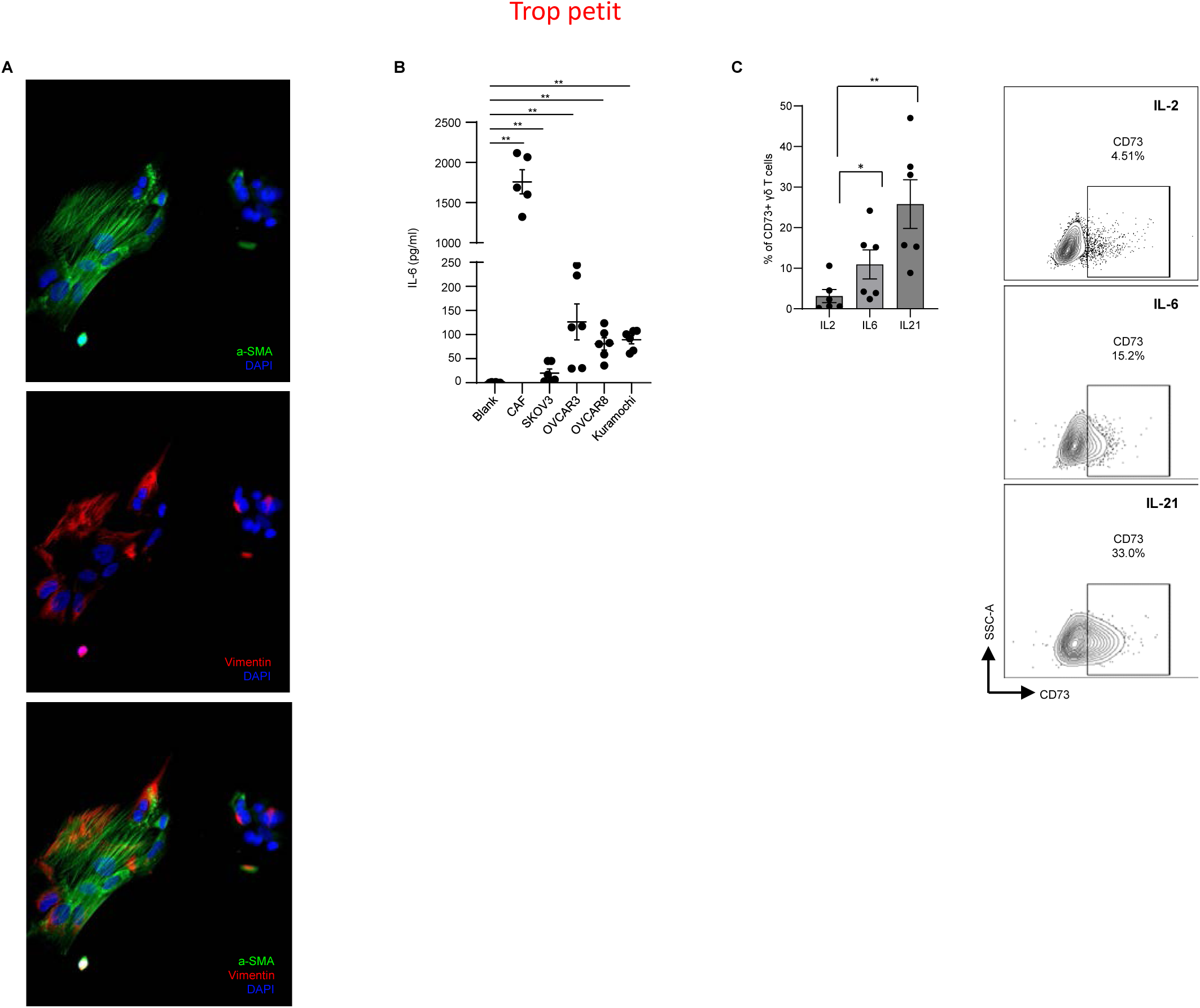
Impact of IL-6 on CD73 expression by γδ T cells. (A) Immunostaining of CAFs with anti-α-SMA and anti-vimentin antibodies, followed by DAPI. (B) IL-6 production was measured in the supernatants of CAFs and four ovarian cell lines (SKOV3, OVSCAR3, OVCAR8 and Kuramochi). Cells were seeded at 1.10^5^ cells/500µl and medium was replaced after 24 hours. Supernatants were harvested after an additional 24 hours and IL-6 was measured by ELISA. (C) After purification, γδ T cells were activated and amplified in the presence of IL-2 alone, and IL-2 with IL-6 or IL-21. CD73 expression was analysed by flow cytometry after seven days. The left panel shows cumulative data from six patients, while the right panel displays a representative dot plot from a single sample. ns: not significant, *: p<0.05, ****: p<0.0001

## Discussion

Investigating the presence, functions and location of immune cells in the TME is essential for developing effective immunotherapy strategies. In this study, we report the presence of CD73+ γδ T cells, a regulatory γδ T-cell subset with immunosuppressive functions, in the ovarian TME of HGSOC and show that their density is associated with poorer clinical outcome in these patients (Fig 1). Although γδ T cells have often been associated with good cancer prognosis (33), several reports in ovarian cancer have provided evidence of their negative impact. For instance, γδ T cells have been positively associated with advanced clinicopathological features of ovarian cancer (34). Moreover, the presence of Vδ1+ cells correlates with progression-free and OS in patients with triple-negative breast cancer (35). Conversely, CD73+ γδ T cell infiltration has been correlated with worse clinical outcome in patients with breast cancer (31). In line with our findings in ovarian cancer, the CD73+ γδ T-cell subset appears to be involved in poor outcome in some cancer types, but the involved mechanisms remain to be elucidated.

By combining scRNA-seq, high-plex IMC and *ex vivo* flow cytometry analysis of freshly resected ovarian tumors, we described in detail the phenotype and ecosystem of CD73+ and CD73-γδ T cells. CD73+ γδ T cells produce IL-10 and IL-8 and also express the immune checkpoints PD-L1 and PD-L2, cytokines and ligands that may explain their regulatory functions and may be involved in the immunosuppressive or pro-tumor mechanisms leading to tumor progression. Indeed, IL-10 impairs the functions of effector cells, such as T and natural killer cells, while PD-L1 and PD-L2 contribute to tumor immune evasion by binding to PD-1 and impair the function of all PD-1+ effector immune cells. IL-8, also known as CXCL8, was originally classified as a chemoattractant for neutrophils and myeloid-derived suppressor cells. It is now known to play other important biological activities in various solid cancer types, such as induction/regulation of angiogenesis and promotion of cancer cell growth, survival and migration (36,37). Several studies have shown that IL-8 induces EMT in ovarian cancer cells and favors migration, invasion and metastasis formation (38–40). IL-8 also promotes ovarian cancer growth in 3D spheroid models (41). Moreover, CAF-derived IL-8 promotes the stemness and malignant proliferation of ovarian cancer cells through NOTCH-3 signaling pathways (42). Conversely, CD73-γδ T cells produce IFN-γ, express the PD-1 and TIGIT checkpoint molecules, and also HLA-DR. Several studies have shown that human γδ T cells can express MHC class II molecules and acquire professional antigen-presenting cell characteristics. Thus, ovarian CD73-γδ TILS have effector T cell properties, whereas CD73+ γδ TILS have regulatory T cell properties.

We also performed a spatial analysis of the CD73+ and CD73-γδ T cell ecosystems in ovarian cancer samples. We found that CD73+ γδ T cells were in close contact with tumor cells expressing more mesenchymal markers, such as ZEB-1, a transcription factor associated with EMT. This is consistent with CD73+ γδ T cell ability to produce IL-8, which has been described as an inducer of several transcription factors involved in EMT and tumor cell migration, such as ZEB-1 and snail (43). In addition, tumor cells close to CD73+ γδ T cells also expressed CD73, which is involved in the generation of the immunosuppressive molecule adenosine from AMP present in the TME. Conversely, tumor cells adjacent to CD73-γδ TILs expressed the immune checkpoint PD-L1 that can be induced by IFN-γ, a cytokine largely produced by CD73-γδ T cells ((44) and our experiments). The spatial analysis also revealed that fibroblasts/CAFs in contact with CD73+ γδ T cells are functionally distinct from those in contact with the CD73-subset. Specifically, one of the CAF clusters in contact with the CD73+ γδ T-cell subset expressed CD73. Several studies have described CD73+ CAF subsets as tumor-promoting or immunosuppressive CAFs. Using a 41-plex antibody panel designed to discriminate the different CAF phenotypes by mass cytometry, Bodenmiller’s group identified a subset, which they termed tumor-like CAFs (tCAFs), with tumor-promoting and immunosuppressive effects and characterized by CD73 and CD10 expression (45). Another study using a mouse model of soft tissue sarcoma identified a CAF subset with a glycolytic signature and characterized by CD73 expression. These glycolytic CAFs rely on GLUT-1-dependent expression of CXCL16 to prevent cytotoxic T-cell infiltration into the tumor. Therefore, CD73+ CAFs could be considered as immunosuppressive or pro-tumor CAFs. In the T-cell compartment, the spatial analysis revealed a cluster of CD4⁺ T cells expressing HLA-DR and/or PD-1 in proximity to the CD73⁻ γδ T-cell subset, suggesting a population with features consistent with recent activation or immune engagement. Conversely, those in contact with CD73+ γδ T cells expressed CD73, PD-1, CD39 and LAG-3, suggesting an exhausted phenotype. The spatial analysis of CD8+ T cells highlighted a different clustering between cells in contact with CD73- and CD73+ γδ T cells, particularly concerning the expression of immune checkpoint factors. Indeed, those in contact with CD73+ γδ T cells expressed PD-1 and CD39, whereas those contact with CD73+ γδ T cells expressed CD73 and Lag-3.

Cytokines can promote or inhibit tumor cell differentiation and immune cell differentiation/functions and angiogenesis, and ultimately affect the patients’ response to therapy (8). Ovarian tumors are characterized by abnormal expression of many cytokines, such as IL-6, IL-8 and IL-10, that affects all TME components and also patient survival (29). IL-8 and IL-10 are produced by CD73+ γδ T cells and IL-6 by cells in their close environment. In addition, γδ T-cell subsets can also produce cytokines with immunosuppressive properties, such as IL-17A and TGF-β and can be involved in the immunosuppressive mechanisms of γδ T cells and/or in their effect on tumor growth, as demonstrated in human colorectal cancer (18). However, in the present study, ovarian cancer CD73+ γδ T cell subsets did not produce these two cytokines (data not shown).

Studies in other cancer types demonstrated the existence of immunosuppressive or pro-tumor γδ T-cell subsets. Daley et al showed in a murine pancreatic cancer model that tumor-infiltrating γδ T cells have potent immunosuppressive functions through the PD-1/PD-L1 axis (20). Specifically, 50% of infiltrating γδ T cells express PD-L1 and induce αβ T-cell exhaustion through this mechanism. In our study, regulatory γδ T cells were characterized by the expression of CD73 and of various immune checkpoint factors, such as PD-L1. These findings suggest that, beyond the immunosuppressive effects mediated by IL-10 and IL-8, CD73⁺ γδ T cells may further promote immunosuppression and tumor progression via immune checkpoint factor expression and adenosine production. In colorectal cancer, Lynch’s group described a γδ T-cell subset that expresses NKp80, produces AREG and induces tumor cell proliferation (46). The existence of pro-tumor and immunosuppressive γδ T-cell subsets is now well established and their role in some cancer types has been demonstrated; however, they represent a heterogeneous population with different mechanisms.

Chronic inflammation is a cancer hallmark and is involved in tumor development and progression. Inflammatory cytokines mediate chronic inflammation and are involved in cancer progression by regulating, for example, immune cell functions. IL-6, which is present at high levels in the ovarian TME, is associated with poor prognosis and increased tumor progression. We and others reported that in ovarian tumors, IL-6 is mainly produced by CAFs and induces CD73 expression by γδ T cells (Fig. 7 and 31). Thus, the proximity between CD73+ γδ T cells and CAFs, observed by IMC, suggests crosstalk mechanisms whereby CAF-produced IL-6 would lead to the recruitment or polarization of γδ T cells into regulatory CD73+ γδ T cells. By a positive feedback, adenosine produced by CD73+ cells would increase IL-6 production in the ovarian TME (Fig. 8). These findings are consistent with the work by Hu et al, who showed that in human breast cancer, CAF-derived IL-6 induces γδ T-cell differentiation into the regulatory CD73+ subset which produces high levels of adenosine following the activation of the IL-6/IL6-R/STAT3 signaling pathway (31). They also showed that CD73+ γδ T cells could in turn promote IL-6 secretion by CAFs through activation of the adenosine/A2BR/p38 MAPK pathway, thereby forming a positive feedback loop (31). This interplay between CAFs and CD73+ γδ T cells could be a general mechanism in solid tumors. The role of IL-8 in EMT and tumor cell migration cells could also be included in this regulation loop, as shown in Figure 8. Consistently, our transcriptomic analysis of ovarian tumors revealed that high expression levels of CD73 were associated with increased expression of TRDC, IL6, IL10, and IL8, further supporting the existence of this immunoregulatory and pro-tumoral network in the TME (Fig 2). Preliminary data also indicate the presence of CD73⁺ γδ T cells in pancreatic and colorectal cancers (data not shown), however, their associated cellular ecosystem remains to be investigated. Overall, our data provide a better understanding of the role of regulatory γδ T cells in the tumor microenvironment and allow to propose mechanisms involving γδ T cells in the creation of a specific immunosuppressive environment (Fig. 8), which could be a more general and existing mechanism in solid tumors with a poor evolution.

**Figure 8.**
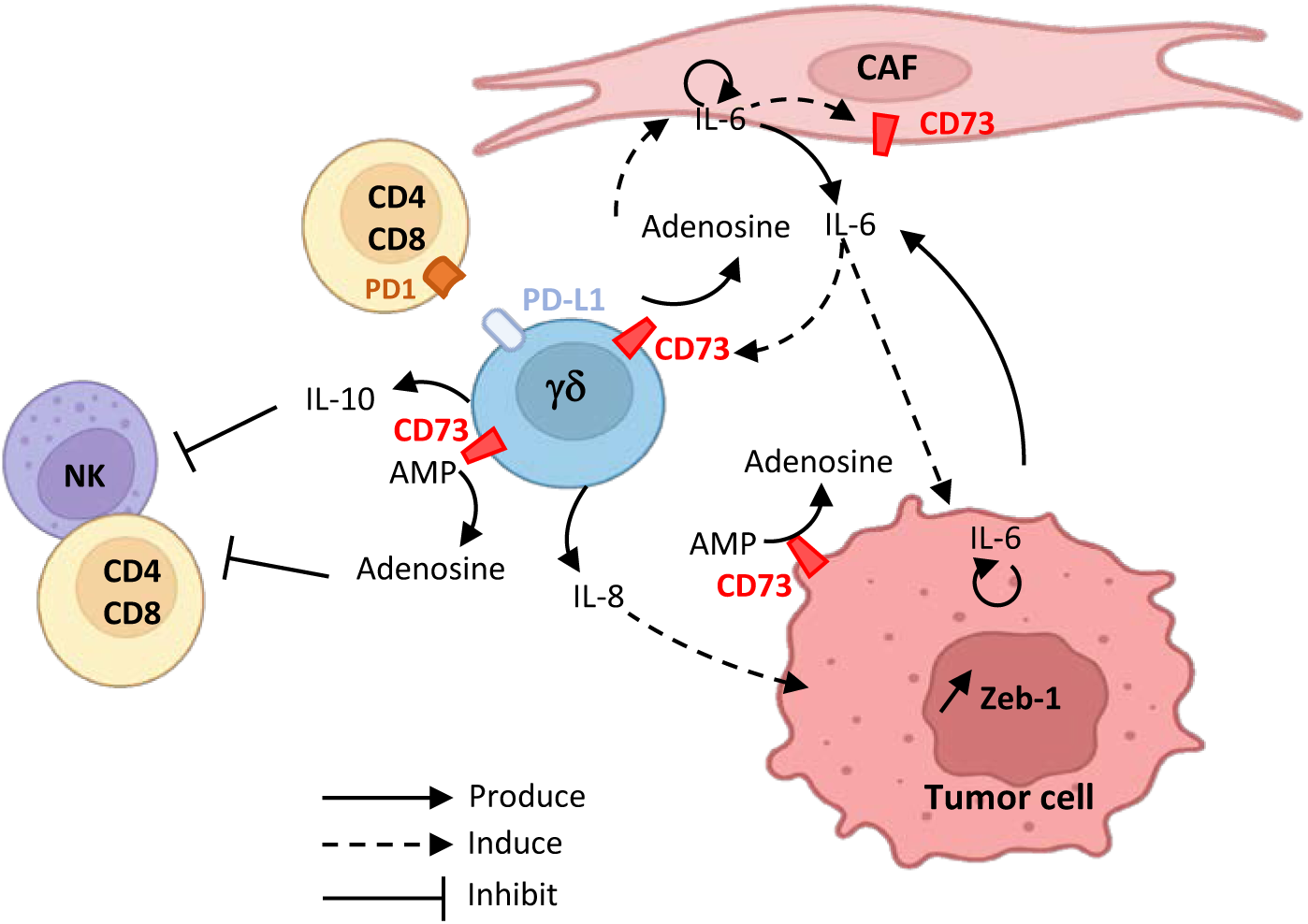
Schematic description of immunosuppressive mechanisms in the neighborhood of CD73+ γδ T cells.

## Acknowledgements

This work was supported by the Institute National de la Santé et de la Recherche Médicale (INSERM); Université Montpellier; the Institut Régional du Cancer de Montpellier (ICM), the SIRIC Montpellier Cancer (Grant INCa_Inserm_DGOS_12553), the Fondation ARC and the Ligue contre le Cancer. Ghita Chabab is supported by a Ph. D. studentship from the Fondation pour la Recherche Médicale and Fondation ARC. We acknowledge the Clinical Resources Center of the Montpellier Cancer Institute (CRB-ICM, n°BB-033-00059). We acknowledge the imaging facility MRI, member of the national infrastructure France-BioImaging infrastructure supported by the French National Research Agency (ANR-10-INBS-04, Investments for the future).

