## Supplementary tables and figures for "Deciphering the CD73⁺ Regulatory γδ T Cell ecosystem associated with poor survival in Ovarian cancer"

| Parameter | Long-term survivors (n=48) | Short-term survivors (n=43) | Total (n=91) | p-value |
|---|---|---|---|---|
| Median age at diagnosis (range) | 62 (22-78) | 63 (33-76) | 62 (22-78) | 0.51 (Mann-Whitney) |
| Neoadjuvant treatment* | | | | |
| Yes | 8 | 6 | 14 | 0.72 (Chi2) |
| No | 40 | 37 | 77 | |
| FIGO stage | | | | |
| III | 42 | 37 | 79 | 0.84 (Chi2) |
| IV | 6 | 6 | 12 | |
| Location | | | | |
| Peritoneum | 13 | 12 | 25 | |
| Ovary | 34 | 27 | 61 | 0.32 (Fisher) |
| Other | 1 | 4 | 5 | |
| Lymph node invasion | | | | |
| N- | 12 | 8 | 20 | 0.37 (Chi2) |
| N+ | 32 | 34 | 66 | |
| Missing | 4 | 1 | 5 | |

*all samples used in this study were from biospies at diagnosis and are, thus, chemonaïve

**Table S1. Clinicopathological characteristics of ovarian cancer cohort**

A

| Target | Clone | Supplier |
|---|---|---|
| CD45 | HI30 | BD Biosciences |
| CD3 | SK7 | BD Biosciences |
| GD | IMMU510 | Beckman Coulter |
| CD73 | AD2 | BD Biosciences |
| CD39 | TU66 | BD Biosciences |
| PD1 | EH12.1 | BD Biosciences |
| PD-L1 | MIH1 | BD Biosciences |
| PD-L2 | MIH18 | BD Biosciences |
| TIGIT | 741182 | BD Biosciences |
| TIM3 | 7D3 | BD Biosciences |
| Viability dye | - | Beckman Coulter |

B

| Target | Clone | Supplier |
|---|---|---|
| CD45 | HI30 | BD Biosciences |
| CD3 | SK7 | BD Biosciences |
| GD | IMMU510 | Beckman Coulter |
| CD73 | AD2 | BD Biosciences |
| IL10 | JES3-9D7 | BD Biosciences |
| IL8 | G265-8 | BD Biosciences |
| IFN-$\gamma$ | B27 | BD Biosciences |
| Viability dye | - | Beckman Coulter |

**Table S2. Antibodies used for flow cytometry experiments.**
Antibodies used for phenotyping are listed in table (A). Antibodies used for cytokine measurment are listed in table (B).

| Target | Clone | Supplier | Tag |
|---|---|---|---|
| Pan CK | AE1/AE3 | SantaCruz | 148 Nd |
| Ecadherin | 24E10 | StandardBiotools | 158 Gd |
| Alpha-SMA | 1A4 | StandardBiotools | 141 Pr |
| Vimentin | D21H3 | StandardBiotools | 143 Nd |
| CD31 | EPR3094 | StandardBiotools | 151 Eu |
| Zeb1/2 | 2A8A6 | Novus | 170 Er |
| CD45 | D9M8I | CellSignaling tech | 89Y |
| CD20 | EP459Y | Abcam | 142 Nd |
| CD3 | D7A6E | CellSignaling tech | 152Sm |
| CD4 | EPR6855 | StandardBiotools | 156 Gd |
| Gamma Delta | H-41 | CellSignaling tech | 161 Dy |
| CD8a | C8/144B | StandardBiotools | 162 Dy |
| FoxP3 | 263A/E7 | StandardBiotools | 155 Gd |
| CD14 | EPR3653 | StandardBiotools | 144 Nd |
| CD16 | EPR16784 | StandardBiotools | 146 Nd |
| CD11b | EPR1344 | StandardBiotools | 149 Sm |
| CD11c | EP1347Y | Abcam | 154 Sm |
| CD68 | KP1 | StandardBiotools | 159 Tb |
| CD66b | REA306 | Miltenyi | 153 Eu |
| CD15 | MMA | Abcam | 163 Dy |
| CD163 | EDHu-1 | StandardBiotools | 147 Sm |
| CD208 | REA295 | Miltenyi | 160 Gd |
| CD57 | NK1 | Abcam | 145 Nd |
| HLA-DR | TAL-1B5 | Abcam | 174 Yb |
| Granzyme-B | EPR20129-217 | StandardBiotools | 167 Er |
| Ki-67 | B56 | StandardBiotools | 168 Er |
| CD73 | D7F9A | CellSignaling tech | 166 Er |
| CD39 | EPR20627 | Abcam | 176 Yb |
| Lag3 | D2G4O | CellSignaling tech | 164 Dy |
| PD-1 | EH33 | CellSignaling tech | 165 Ho |
| CD366/TIM-3 | D5D5R | CellSignaling tech | 171 Er |
| PD-L1 | SP142 | Abcam | Uncoupled |
| aRb | Polyclonal | StandardBiotools | 175 Lu |

**Table S3. Antibodies used for imaging mass cytometry**

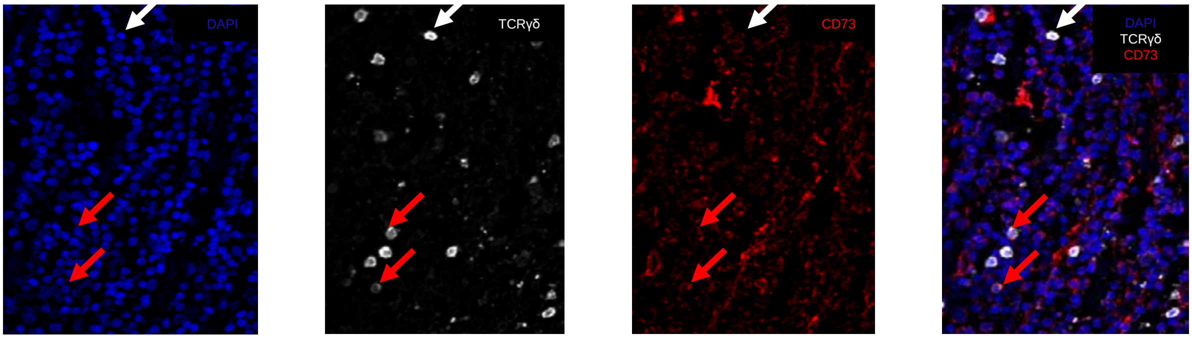


**Supplementary Fig.1: Immunofluorescence on ovarian tumors.**

Detection of γδ T cells in 91 ovarian tumor samples by immunofluorescence. The figure shows the staining of one representative tumor. The red arrow indicates CD73+ γδ T cells and the white arrow, the CD73- γδ T cells.

A

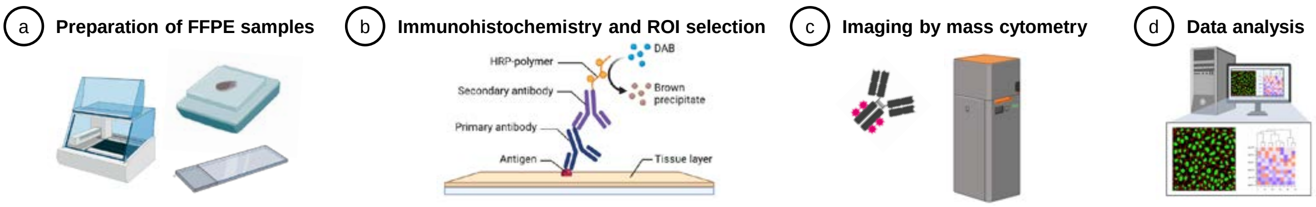


B

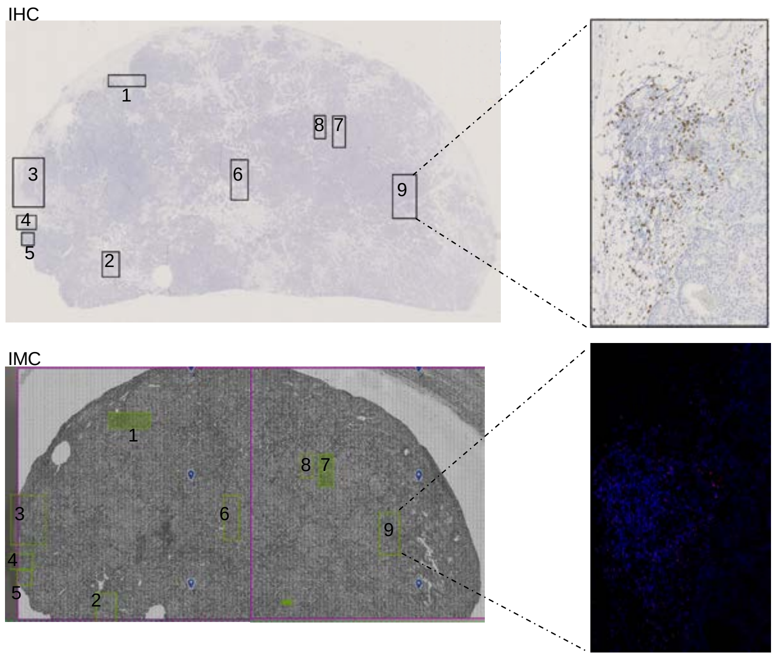


**Supplementary Fig.2: Staining procedures and ROI selection**

A) Staining procedures

(a) 5μm formalin-fixed paraffin-embedded (FFPE) tissue sections are deparaffinized. Two serial sections are prepared, one for IHC and one for IMC staining;

(b) IHC sections are stained with the H-41 antibody to detect γδ T cells. ROIs are selected based on the presence of γδ T cells;

(c) IMC sections are stained with the H-41 antibody to detect γδ T cells. ROIs are selected based on the presence of γδ T cells;

(d) Data scaling and normalization, statistical tests, and neighborhood analysis.

B) ROI selection

After IHC, regions rich in γδ T cells are selected for ablation and IMC analysis

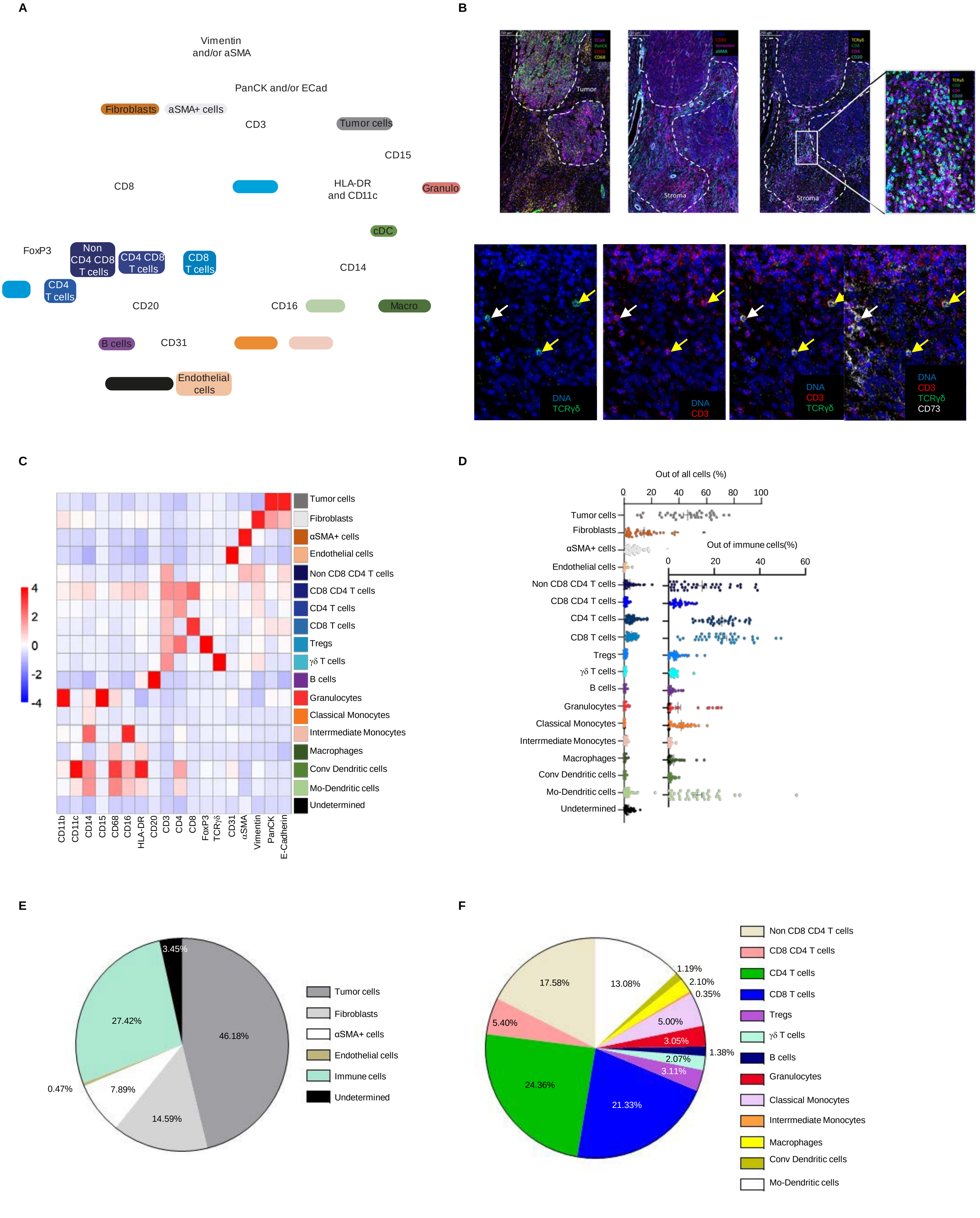


**Supplementary Fig.3: Description of ovarian tumor tissue description by imaging mass cytometry analysis**

(A) Gating strategy used for cell type identification. (B) Examples of IMC staining. In the lower panel, CD73- γδ T are indicated by the yellow arrow and CD73+ γδ T by the white arrow (C) Heat map showing the mean expression of all markers in the identified cell type in ovarian tumor tissue samples. (D) Dot plot showing the frequency of each cell type in each ROI. (e) Pie chart showing the mean proportion of each identified cell type in ROIs. (f) Pie chart showing the mean value of each identified immune cell type in ROIs.

Tregs, Regulatory T cells; cDC, conventional Dendritic Cells; Mo-DC, Monocyte-derived Dendritic cells ; Cl-MONO, classical monocytes; Int-Mono, intermediate monocytes, Macro, macrophages, Granulo, granulocytes; neutro, neutrophiles.

A

| Cluster | 1 | 2 | 3 | 4 | 5 | 6 | 7 | 8 |
|---|---|---|---|---|---|---|---|---|
| **CD73- γδ T cell** | 17.18 | 9.80 | 5.95 | 3.53 | 22.64 | 18.67 | 3.91 | 18.30 |
| **CD73+ γδ T cell** | 10.49 | 10.49 | 18.16 | 2.06 | 13.86 | 6.37 | 32.40 | 6.18 |

B

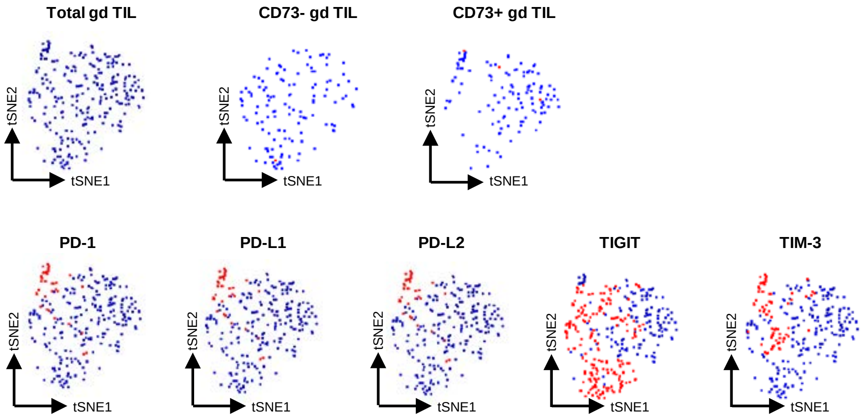


| Marker | PD-1 | PD-L1 | PD-L2 | TIGIT | TIM-3 |
|---|---|---|---|---|---|
| **CD73- γδ T cell** | 3.72 | 30.1 | 8.78 | 54.4 | 13.9 |
| **CD73+ γδ T cell** | 19.5 | 50.8 | 35.2 | 31.2 | 24.2 |

C

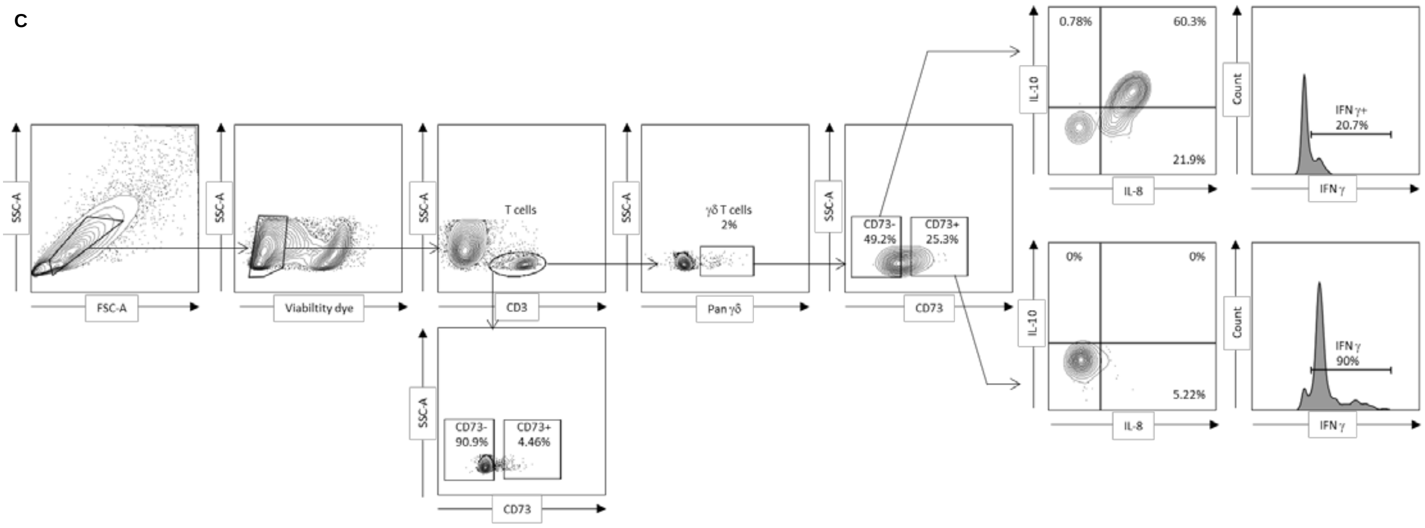


**Supplementary Fig.4: Phenotype of γδ T cell subsets.**

(A) IMC analysis: Frequency of the CD73- and CD73+ γδ T-cell subsets in the identified clusters based on the clustering of Fig. 2f. (B) Upper panels: tSNE plots showing the clustering of total, CD73+ and CD73- γδ T cells from fresh ovarian tumor samples (n=7) analyzed by flow cytometry. Lower panels: tSNE plots showing the expression of each indicated protein in the total γδ T cell clusters; the table shows the percentage of positive CD73+ and CD73- γδ T cells. (C) Representative flow cytometry gating, dot plots and histograms showing the pourcentage of CD73- and CD73+ γδ T cells expressing IL10, IL8 and IFNγ.

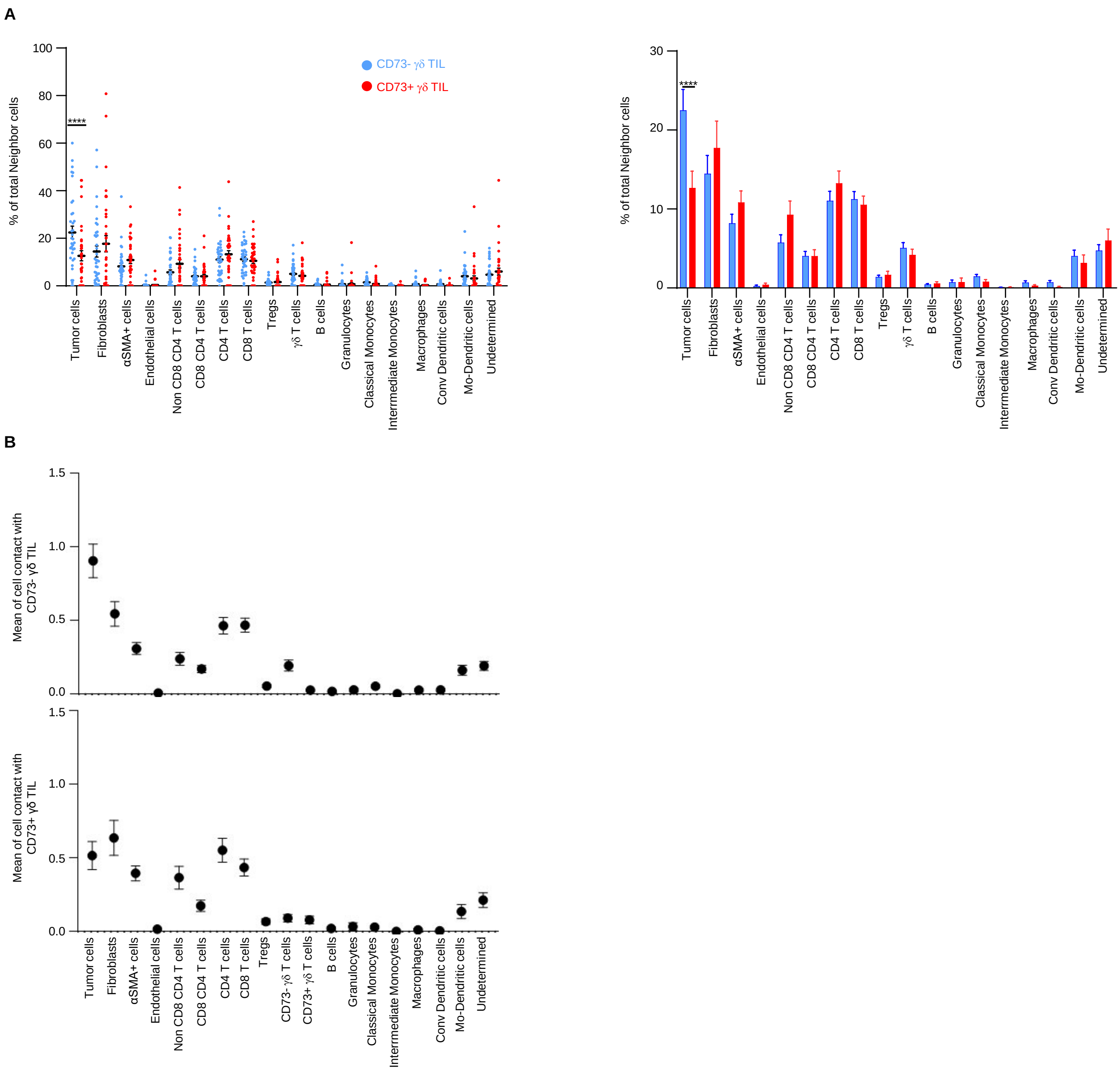


**Supplementary Fig. 5: Characterization of neighbor cells of CD73+ and CD73- γδ T cell subsets.**

(A) Each circle represents the percentage of neighbors for the indicated cell type of CD73+ (red) and CD73- (blue) γδ T cells in a ROI; ****p <0.0001 (Wilcoxon signed rank test). (B) mean of cells in contact with CD73+ (lower panel) and CD73- (upper panel) γδ T cells.

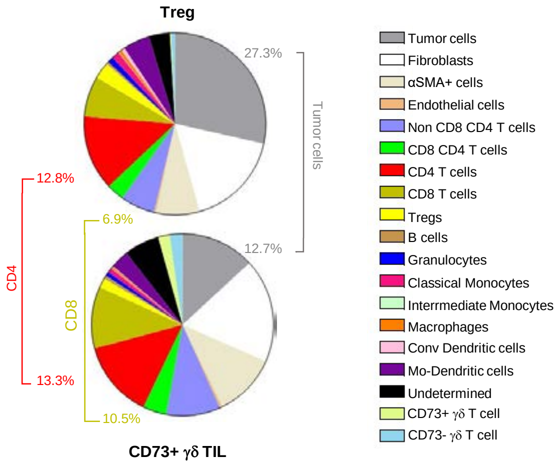


**Supplementary Fig. 6: Comparison of CD4 regulatory T cells and CD73+ γδ T cells neighborhoods**

Pie chart showing the mean value of each identified cell type in the selected ROIs. Upper panel: Neighbors of Treg cells; Lower panel: Neighbors of CD73+ γδ T cells.

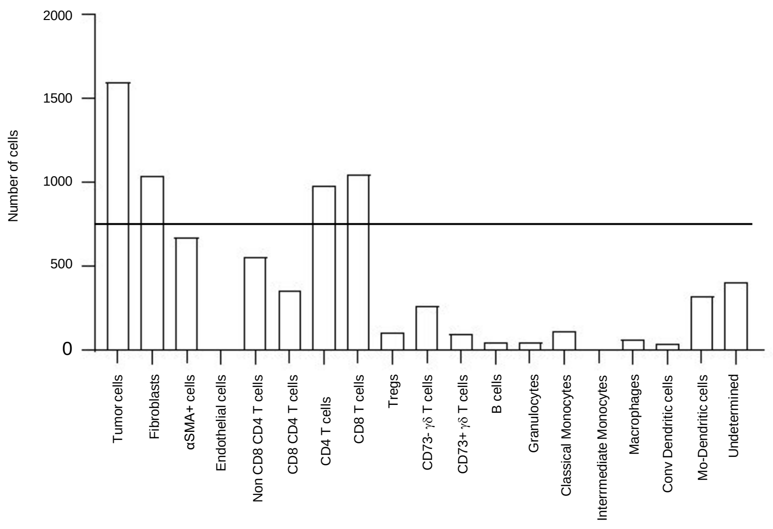


**Supplementary Fig. 7: Number of cells per cell type in the neighborhood of γδ T cells**

**A**

| **Tumor cells (clusters)** | **1** | **2** | **3** | **4** | **5** | **6** | **7** | **8** |
|---|---|---|---|---|---|---|---|---|
| Neighbors of CD73- γδ T cells | 25.60 | 8.09 | 16.27 | 7.25 | 14.26 | 14.88 | 10.80 | 2.85 |
| Neighbors of CD73+ γδ T cells | 20.10 | 6.88 | 8.99 | 22.75 | 17.46 | 14.28 | 5.29 | 4.23 |

**B**

| **Fibroblasts (cluster)** | **1** | **2** | **3** | **4** | **5** | **6** | **7** |
|---|---|---|---|---|---|---|---|
| Neighbors of CD73- γδ T cells | 24.35 | 19.12 | 2,75 | 14.30 | 18.98 | 12.38 | 8.11 |
| Neighbors of CD73+ γδ T cells | 14.47 | 13.50 | 6.11 | 32.80 | 24.11 | 5.47 | 3.54 |

**C**

| **CD4 T cells (cluster)** | **1** | **2** | **3** | **4** | **5** | **6** | **7** | **8** |
|---|---|---|---|---|---|---|---|---|
| Neighbors of CD73- γδ T cells | 15.32 | 5.78 | 13.44 | 29.05 | 13.15 | 1.30 | 9.83 | 12.14 |
| Neighbors of CD73+ γδ T cells | 7.34 | 6.64 | 1.75 | 23.43 | 10.14 | 2.10 | 38.81 | 9.79 |

**D**

| **CD8 T cells (cluster)** | **1** | **2** | **3** | **4** | **5** | **6** | **7** | **8** | **9** | **10** |
|---|---|---|---|---|---|---|---|---|---|---|
| Neighbors of CD73- γδ T cells | 2.46 | 11.55 | 5.16 | 19.53 | 24.57 | 13.03 | 9.58 | 4.54 | 1.10 | 8.48 |
| Neighbors of CD73+ γδ T cells | 2.31 | 8.33 | 3.70 | 10.18 | 24.54 | 12.04 | 3.70 | 25.00 | 2.78 | 7.41 |

**Supplementary Fig. 8. Percentage of clusters in the neighborhood of CD73+ and CD73- γδ T cell subsets for each cell type**

(A) Tumor cell clusters; (B) Fibroblast clusters; (C) CD4+ T-cell clusters; (D) CD8+ T-cell clusters.

A

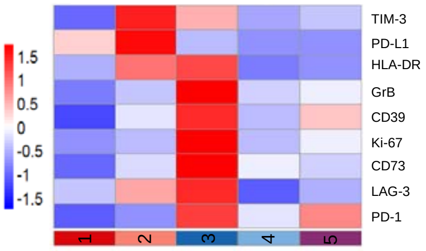


| Cluster | 1 | 2 | 3 | 4 | 5 |
|---|---|---|---|---|---|
| Neighbor of CD73- γδ T cell | 15.21 | 23.57 | 25.10 | 18.63 | 17.49 |
| Neighbor of CD73+ γδ T cell | 3.84 | 17.31 | 55.77 | 12.50 | 10.58 |

B

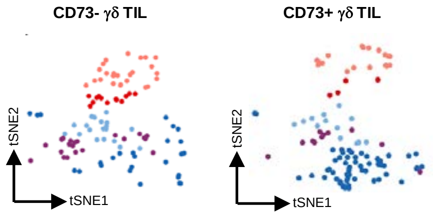


C

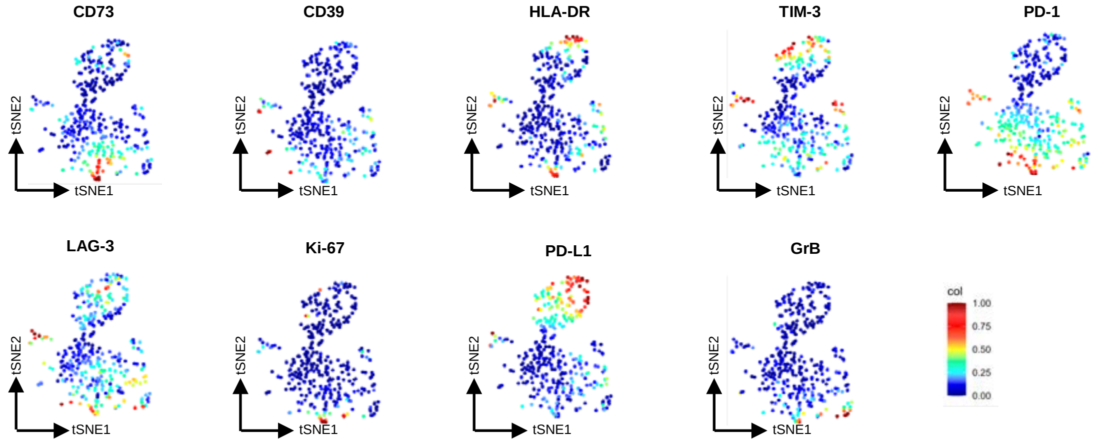


**Supplementary Fig. 9. Phenotype of γδ T cells present in the neighborhood of CD73+ and CD73- γδ T cells**

(A) Heat map showing the clustering of γδ T cells based on the expression of the listed markers (left panel) and table showing the percentage of each cluster (right panel). (B) tSNE plots showing cluster repartition in CD73+ and CD73- γδ T cells. (C) tSNE plots showing the expression of each analyzed marker in the total γδ T cell clusters.
